# *In vivo* marker of brainstem myelin is associated to quantitative sleep parameters in healthy young men

**DOI:** 10.1101/2023.07.17.549285

**Authors:** Puneet Talwar, Michele Deantoni, Maxime Van Egroo, Vincenzo Muto, Daphne Chylinski, Ekaterina Koshmanova, Mathieu Jaspar, Christelle Meyer, Christian Degueldre, Christian Berthomier, André Luxen, Eric Salmon, Fabienne Collette, D.-J. Dijk, Christina Schmidt, Christophe Phillips, Pierre Maquet, Siya Sherif, Gilles Vandewalle

## Abstract

Brain structural integrity has been suggested to contribute to the variability in human sleep quality and composition. The associations between sleep parameters and the regional integrity of subcortical structures implicated in sleep-wake regulation remain, however, largely unexplored. The present study aimed at assessing association between quantitative Magnetic Resonance Imaging (qMRI)-derived marker of the myelin content of the brainstem with the variability in the sleep electrophysiology in a large sample of healthy young men (N=321;∼22y). Generalized Additive Model for Location, Scale and Shape (GAMLSS) was used to seek associations between sleep metrics and Magnetisation Transfer saturation (MTsat) qMRI values, proxy for myelin content. Separate GAMLSS revealed that sleep onset latency and slow wave sleep intensity were significantly associated with MTsat-derived myelin estimates in the brainstem (p_corrected_≤.03), with overall higher MTsat value associated with values reflecting better sleep quality. The association changed with age, however (MTsat-by-age interaction - p_corrected_≤.03), with higher MTsat value linked to better values in the two sleep metrics in the individuals of our sample aged ∼18 to 20y. Similar associations were detected across different parts of the brainstem (p_corrected_≤.03), suggesting that the overall maturation and integrity of the brainstem was associated with both sleep metrics. Our results suggest that myelination of the many reticular nuclei of the brainstem essential to regulation of sleep is associated with inter-individual differences in sleep characteristics during early adulthood. They may have implications for sleep disorders or neurological diseases related to myelin.

## Introduction

Sleep is essential to physical and mental health. Poor sleep is a predictor of mental health disorders, accelerated cognitive aging and neurodegeneration (Zeitzer, 2013). Although sleep undergoes profound changes over the lifetime (Carrier et al., 2011), it is remarkably stable within an individual over a shorter life period (e.g. a few months/years) (Tucker et al., 2007). The architecture of sleep is, however, highly variable across the individuals, even when they are very healthy (Tucker et al., 2007). A better understanding of such variability could provide important keys on the brain bases of a better sleep and for individually tailored sleep interventions.

The regulation of sleep heavily relies on nuclei of the reticular formation in the brainstem. They interact with each other and with nuclei of the diencephalon and basal forebrain to set vigilance state over the cortex (Zeitzer, 2013, Scammell et al., 2017). Monoaminergic neurons, including the noradrenergic projection originating in the locus coeruleus (LC) and the serotonergic projection from the dorsal raphe nucleus, together with the cholinergic projection from laterodorsal tegmental nucleus (LDT) and the pedunculopontine tegmental (PPT) nucleus, project to most cortical territories to promote wakefulness, when more active, or allow sleep when less active (Scammell et al., 2017). LC, LDT and PPT are also central to the switch between Rapid Eye Movement Sleep (REMS) and Slow Wave Sleep (SWS) as well as the oscillatory modes most typical of REMS and SWS (Kayama and Koyama, 2003, Anaclet and Fuller, 2017, Benarroch, 2018). The norepinephrine (NE)-LC system also contributes to the organisation of sleep oscillations including slow wave and spindles (Osorio-Forero et al., 2021).

Most of the knowledge about the involvement of subcortical nuclei in sleep and wakefulness regulation arises from animal studies that used lesions, pharmacology or conditional stimulation to demonstrate the essential role of a given subcortical nucleus. Whether the natural variability in the structure or functioning of these nuclei contributes to the variability in observed sleep phenotypes remains, however, mostly unknown. This is also true in humans where, aside from rare studies linking the integrity of the LC to subjective reports of quality of sleep (Van Egroo et al., 2021, Van Egroo et al., 2022), *in vivo* studies of the association between the integrity of the brainstem and sleep are inexistent.

The integrity or the thickness of several cortical areas has been associated with variability in the electrophysiology of sleep of healthy younger and older individuals (Dube et al., 2015, Van Egroo et al., 2019). Many of these associations arise arguably from changes in myelination, which is critical for brain function, including during sleep (Fitzroy et al., 2021). The progressive myelination of the brain occurs throughout adolescence [early (11-14y), mid (15-17y) and late (18-24y)] and continues into early adulthood (22-30y). It results in a substantial increase in the brain’s white matter volume (WMV) over these periods of life. In the central nervous system, myelin is present not only in white matter, but also in varying amounts in many grey matter areas where it is present around neurites near the neuron cell body (Arain et al., 2013, Jamieson et al., 2020b). Most brainstem nuclei constitute densely myelinated regions which undergo progressive myelination until early adulthood (Bouhrara et al., 2021). How this progressive change is reflected in sleep has not been investigated.

Here, we reasoned that the myelination of the nuclei of the brainstem could drive, to some extent, sleep electrophysiology. We characterized a proxy of myelin content over the entire brain using quantitative Magnetic Resonance Imaging (qMRI) in a large sample of healthy young men devoid of sleep disorders. We then extracted myelin proxy values over the brainstem monoaminergic grey matter (bmGM) compartment comprising the LC, raphe, LDT and PPT nuclei. We recorded sleep of all participants in the laboratory using electroencephalography (EEG) and extracted some of the most prominent sleep features related to sleep initiation, sleep continuity, global sleep architecture, as well as markers of the intensity of SWS and REMS. Since the 18-to-31y age range of our sample spanned a critical period for brain maturation and myelination, we further assessed whether potential links between sleep metrics and the myelin marker over bmGM compartment would change with age. Given the scarcity of the available literature and the critical age range of our sample we did not have a priori hypothesis on the direction of the potential links between the sleep metrics and myelin proxy and corrected accordingly for the multiple comparisons.

## Results

Following 3 weeks of stable sleep-wake timing, we recorded sleep of 321 healthy young men (22.1y±2.7) under EEG (**Table 1**). We extracted six sleep metrics: 1) sleep onset latency (SOL), related to the initiation of sleep; 2) Slow wave energy (SWE) during SWS, corresponding to the cumulated overnight EEG power of delta band (.5-4 Hz), an accepted marker of sleep need and SWS intensity (Gillberg and Akerstedt, 1991); 3) sleep efficiency (SE; ratio between sleep time and time in bed), to assess overall sleep quality and continuity (Basiri et al., 2017); 4) REMS percentage, to reflect the global architecture of sleep; 5) the cumulated theta power during REMS, associated with REMS intensity over its most typical oscillatory activity (Riemann et al., 2012) and 6) number of arousals during REMS, to characterise sleep continuity (Riemann et al., 2012). All participants subsequently underwent a qMRI protocol through which we computed the Magnetization Transfer saturation (MTsat) value, which is a semi-quantitative MRI measure linked to the myelin content (Schmierer et al., 2004), for each voxel of the brain prior to extracting the average value of the bmGM compartment. The overview of the study design is provided in **Figure 1**.

**Table 1.** Characteristics of the final participant cohort included in the analyses.

| Characteristic | Mean (SD) |
| --- | --- |
| Sample size (N) | 321 |
| Sex | Men |
| Ethnicity | Caucasian |
| Age (years) | 22.09 (2.74) |
| BMI (kg*m<sup>-2</sup>) | 22.15 (2.43) |
| Anxiety<sup>b</sup> | 2.65 (3.18) |
| Mood<sup>b</sup> | 3.05 (3.48) |
| Sleep Quality<sup>a</sup> | 3.46 (1.79) |
| Daytime sleepiness<sup>a</sup> | 5.96 (3.54) |
| Chronotype<sup>a</sup> | 50.12 (8.25) |
| Total Sleep Time (min) | 451.69 (42.25) |
| N1 duration (min) | 10.77 (8.08) |
| N2 duration (min) | 220.39 (36.77) |
| N3 duration (min) | 99.55 (24.44) |
| Sleep onset latency (min) | 16 (11) |
| Slow wave energy (μV<sup>2</sup>) | 2618214 (2422398) |
| Sleep efficiency, incl. N1 (%) | 93.25 (3.92) |
| REM duration (min) | 120.96 (25.0) |
| REM theta power (4-8Hz) | 103674 (68508) |
| REM arousal (N) | 26 (13) |

Sleep quality was assessed by the Pittsburgh Sleep Quality index (PSQI (Buysse et al., 1989)). Daytime sleepiness was measured by the Epworth Sleepiness Scale (Johns, 1991), Chronotype was assessed by the Morningness-Eveningness Questionnaire (MEQ, (Horne and Ostberg, 1976)). Anxiety was estimated by the Beck Anxiety Inventory (Beck et al., 1993) and Mood was estimated by the 21-item Beck Depression Inventory II (Beck et al., 1988); Total Sleep Time (TST) was extracted from polysomnography recordings.
<sup>a,b</sup> The data was available for 317 and 320 participants respectively.

**Figure 1:**
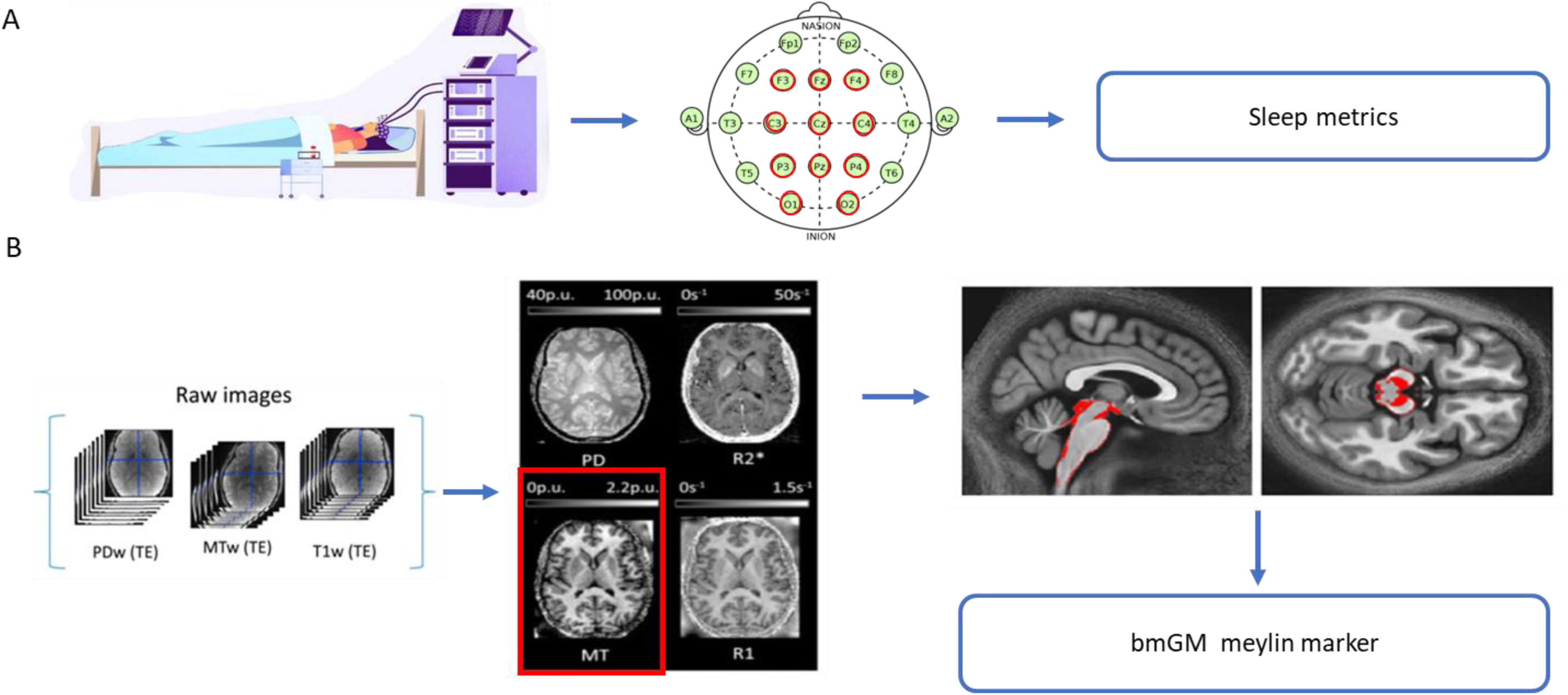
Overview of the study design: **A.** In-lab recordings of habitual sleep to extract sleep macro and microstructure metrics. **B.** 7T MRI MPM acquisitions to generate qMRI maps. MT value (related to myelin content) was averaged was computed over the bmGM compartment.

We first assessed whether our sleep metrics of interest varied with age. Despite the limited age range of our sample, we found significant decreases in SWE (p<0.001), SE (p=0.002), and REM theta power (p=0.001) with age and significant increases in REM arousals (p=0.001), which is in line with previously published age-related changes in these variable (Baker et al., 2016, Sprecher et al., 2016, Li et al., 2018) (**Suppl. Table S1, Suppl. Figure S1**). No significant age-related changes in SOL (p=0.11), and in REM percentage (p = 0.69) were detected. A negative borderline association in brainstem monoaminergic grey matter myelin content was further observed with age (p=0.05) (**Table S1, Suppl. Figure S1**).

### Latency to sleep, SWS intensity and sleep efficiency are associated with the qMRI myelin marker over the monoaminergic brainstem compartment

Given the exploratory nature of our investigation our statistical analyses consisted of GAMLSS which is a flexible distributional regression approach considered as an improvement and extension to the generalized linear models (GLM) and the generalized additive models (GAM) (Marmolejo-Ramos et al., 2023). A first GAMLSS, with SOL as the dependent variable, yielded a significant negative main effect of the MTsat value of the bmGM (**p= 2.2×10^-5^, p_corr_=1.2×10^-4^)** and of age (**p=3.8×10^-6^, p_corr_=2.3×10^-5^)** while controlling for body mass index (BMI), total sleep time (TST), and total intracranial volume (TIV), as well as MRI MPM sequence and scanner (see methods) (**Table 2**, **Fig. 2A**). The same GAMLSS also yielded an interaction between the bmGM MTsat value and age (**p=5.0×10^-5^, p_corr_=0.0003**). To gain insight in this interaction, we split our sample into 3 subsamples of similar size, respectively ranging from 18 to 20y (N=104), from 21 to 23y (N =133) and from 24 to 31y (N=84) (**Suppl. Table S2** for sleep metrics in each subgroup). We then computed GAMLSS for the 3 subgroups separately and found that while the association was significantly negative in the younger subsample (**p=0.006**), i.e. with higher MTsal related to shorter SOL, there was no significant association with the intermediate and the older subsamples (p≥0.36) (**Table 3**, **Fig. 2C**). This suggests that the interaction between MTsat values and age that reflect a change in the slope and/or direction of the association between SOL and MTsat, is mostly driven by the younger individuals of our sample.

**Table 2.** Results derived from GAMLSS when testing for associations between sleep parameters and MTsat values computed over brainstem monoaminergic grey matter (bmGM) with age as interacting variable.

| Sleep parameters | SOL |  | SWE |  | SE |  | REMS % |  | REMS theta Power |  | REMS Arousal (N) |  |
| --- | --- | --- | --- | --- | --- | --- | --- | --- | --- | --- | --- | --- |
|  | Estimate | p-value (95% CI) | Estimate | p-value (95% CI) | Estimate | p-value (95% CI) | Estimate | p-value (95% CI) | Estimate | p-value (95% CI) | Estimate | p-value (95% CI) |
| MTsat value bmGM | -48.60 | **2.2e-05**<br>(-62.1, -35.1) | 41.40 | **0.01** (9.1,73.7) | 0.63 | 0.14<br>(0.34, 0.86) | -0.40 | 0.89<br>(-6.46, 5.67) | 12.1 | 0.28<br>(-9.71, 33.8) | 4.23 | 0.57<br>(-10.60, 19.0) |
| Age | -0.49 | **1.2e-04**<br>(-0.65, -0.34) | 0.38 | **0.04**<br>(0.01, 0.74) | 0.01 | 0.17<br>(0.004, 0.01) | -0.01 | 0.88<br>(-0.07, 0.06) | 0.08 | 0.51<br>(-0.17, 0.33) | 0.07 | 0.41<br>(-0.10, 0.24) |
| Age*MTsat bmGM | 2.11 | **5.1e-05**<br>(1.5, 2.73) | -1.84 | **0.01**<br>(-3.32, -0.37) | -0.03 | 0.12<br>(-0.04, -1.83) | 0.02 | 0.88<br>(-0.26, 0.3) | -0.53 | 0.30<br>(-1.53, 0.47) | -0.15 | 0.65<br>(-0.83, 0.52) |
| BMI | -0.058 | **2.52e-04**<br>(-0.09, -0.027) | -0.034 | 0.12<br>(-0.008, 0.08) | 0.001 | 0.33<br>(-0.001, 0.002) | 0.01 | **0.02**<br>(0.002, 0.02) | 0.03 | **0.03**<br>(0.004, 0.06) | 0.006 | 0.57<br>(-0.03,0.01) |
| TST | 0.002 | **0.03**<br>(0.0002,0.004) | -0.001 | 0.26<br>(-0.004, 0.001) | -4.7e-05 | 0.14<br>(-0.0001, 2.1e-05) | 0.001 | **0.01**<br>(0.0001, 0.001) | 0.001 | 0.23<br>(-0.0005, 0.003) | 0.004 | **2.5e-09**<br>(0.003, 0.005) |
| TIV | 0.001 | 0.11<br>(-0.0001,0.001) | -0.0002 | 0.66<br>(-0.0003, -0.0002) | 8.9e-07 | 0.94<br>(-, -) | -4.4e-05 | 0.60<br>(-0.0002, 0.0001) | -1.2e-05 | 0.97<br>(-3.9e-5, 1.6e-05) | -2.1e-05 | 0.92<br>(-4.4e-05, -2.1e-06) |
| MPM sequence | 0.18 | **0.03**<br>(0.02, 0.34) | -0.14 | 0.23<br>(-0.36, 0.09) | -0.003 | 0.33<br>(-0.01, 0.003) | 0.01 | 0.72<br>(-0.03, 0.05) | 0.01 | 0.94<br>(-0.15, 0.17) | 0.02 | 0.71<br>(-0.09, 0.13) |
| Scanner type | -0.13 | 0.31<br>(-0.37, 0.11) | 0.03 | 0.87<br>(-0.34, 0.39) | -0.0002 | 0.96<br>(-0.01, 0.01) | 0.004 | 0.90<br>(-0.06,0.07) | 0.11 | 0.39<br>(-0.15, 0.36) | -0.01 | 0.91<br>(-0.17, 0.16) |

BMI: body mass index; bmGM: brainstem monoaminergic gray matter compartment; MPM sequence: type of multiparameter sequence, 2 sequence types were used (N = 222 for type 1; N = 99 for type 2- see methods); MRI scanner: data were acquired on 2 separate scanners (N = 286 for scanner 1; N = 35 for scanner 2 – see methods); MTsat: magnetisation transfer saturation; REMS: rapid eye movement sleep; SE: sleep efficiency; SOL: sleep onset latency; SWE: slow wave energy; TIV: total intracranial volume; TST: total sleep time.

**Figure 2:**
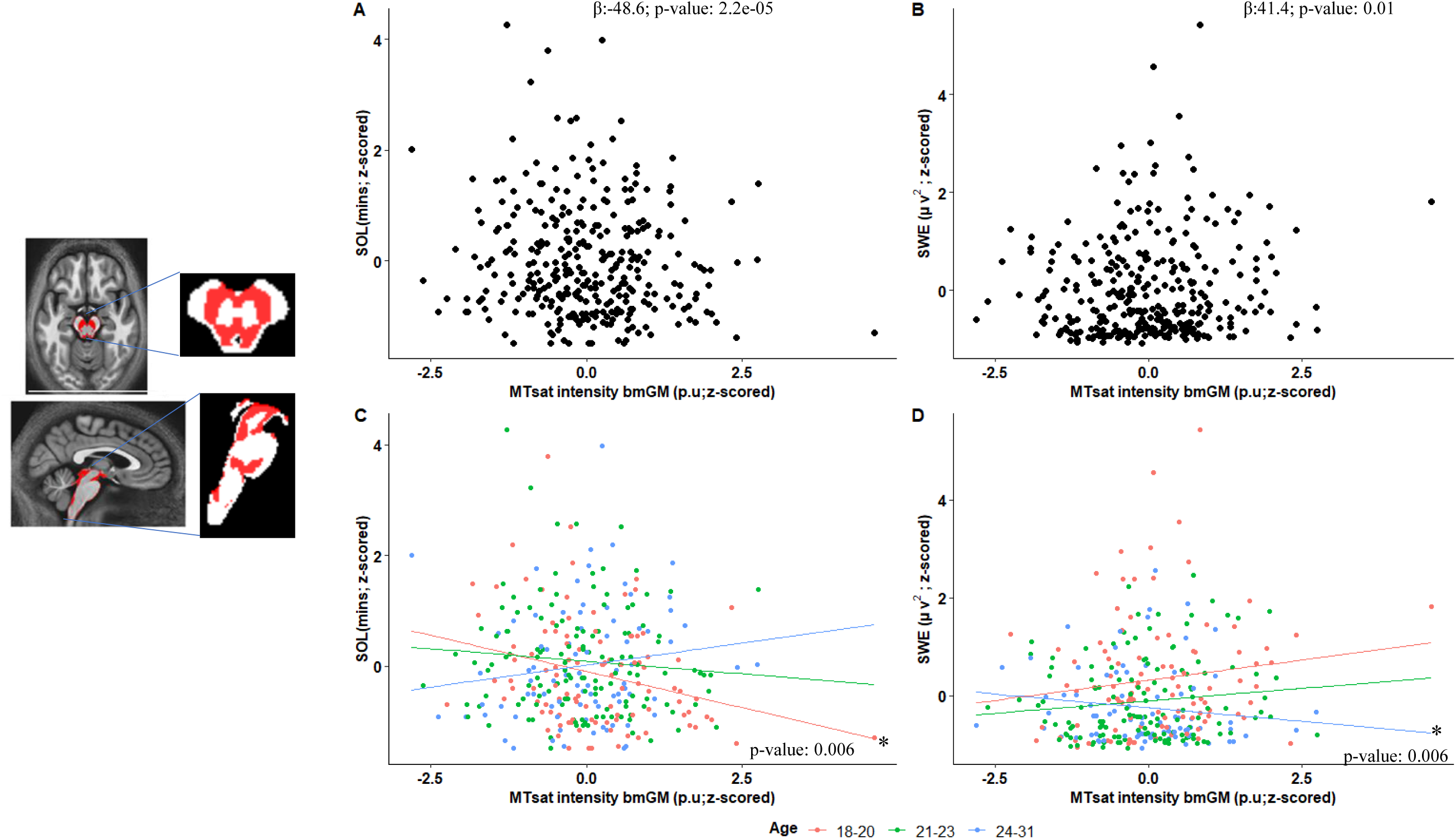
Plots (A-B) depict pearson correlations between sleep onset latency (SOL) and slow wave energy (SWE) with MTsat intensity in bmGM respectively (N=321), see Table 2 for statistical outputs of GAMLSSs. Plots (C-D) shows association between sleep parameters and MTsat values by age group for SOL and SWE with MTsat intensity in brainstem monoaminergic grey matter (bmGM) respectively, see Table 2 and Table 3 for statistical outputs of GAMLSSs.

**Table 3:**
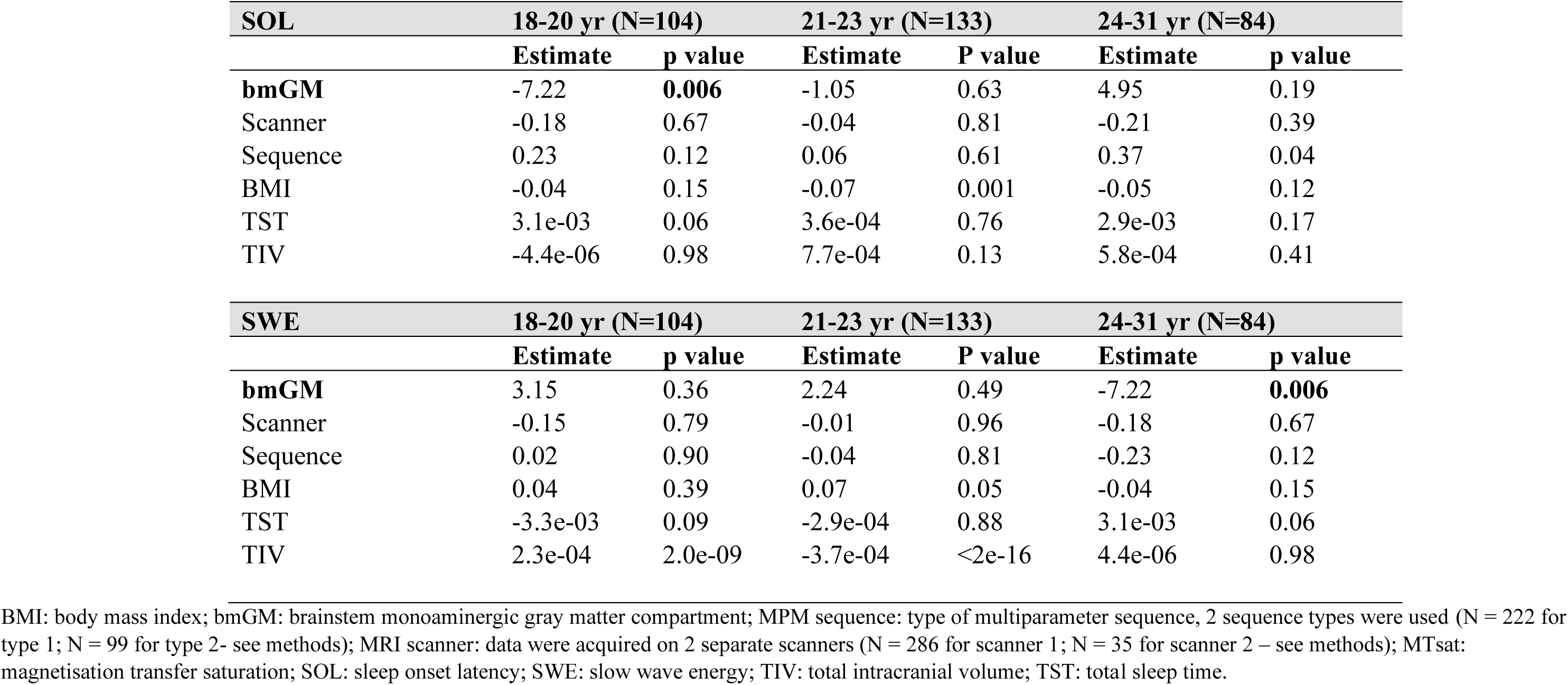
Results derived from GAMLSS in age subgroups when testing for associations between sleep onset latency (SOL), slow wave energy (SWE) or sleep efficiency (SE) and MTsat values computed over the brainstem monoaminergic gray matter compartment (bmGM)

| **SOL** | **18-20 yr (N=104)** |  | **21-23 yr (N=133)** |  | **24-31 yr (N=84)** |  |
| --- | --- | --- | --- | --- | --- | --- |
|  | **Estimate** | **p value** | **Estimate** | **P value** | **Estimate** | **p value** |
| **bmGM** | -7.22 | **0.006** | -1.05 | 0.63 | 4.95 | 0.19 |
| Scanner | -0.18 | 0.67 | -0.04 | 0.81 | -0.21 | 0.39 |
| Sequence | 0.23 | 0.12 | 0.06 | 0.61 | 0.37 | 0.04 |
| BMI | -0.04 | 0.15 | -0.07 | 0.001 | -0.05 | 0.12 |
| TST | 3.1e-03 | 0.06 | 3.6e-04 | 0.76 | 2.9e-03 | 0.17 |
| TIV | -4.4e-06 | 0.98 | 7.7e-04 | 0.13 | 5.8e-04 | 0.41 |

| **SWE** | **18-20 yr (N=104)** |  | **21-23 yr (N=133)** |  | **24-31 yr (N=84)** |  |
| --- | --- | --- | --- | --- | --- | --- |
|  | **Estimate** | **p value** | **Estimate** | **P value** | **Estimate** | **p value** |
| **bmGM** | 3.15 | 0.36 | 2.24 | 0.49 | -7.22 | **0.006** |
| Scanner | -0.15 | 0.79 | -0.01 | 0.96 | -0.18 | 0.67 |
| Sequence | 0.02 | 0.90 | -0.04 | 0.81 | -0.23 | 0.12 |
| BMI | 0.04 | 0.39 | 0.07 | 0.05 | -0.04 | 0.15 |
| TST | -3.3e-03 | 0.09 | -2.9e-04 | 0.88 | 3.1e-03 | 0.06 |
| TIV | 2.3e-04 | 2.0e-09 | -3.7e-04 | <2e-16 | 4.4e-06 | 0.98 |

BMI: body mass index; bmGM: brainstem monoaminergic gray matter compartment; MPM sequence: type of multiparameter sequence, 2 sequence types were used (N = 222 for type 1; N = 99 for type 2- see methods); MRI scanner: data were acquired on 2 separate scanners (N = 286 for scanner 1; N = 35 for scanner 2 – see methods); MTsat: magnetisation transfer saturation; SOL: sleep onset latency; SWE: slow wave energy; TIV: total intracranial volume; TST: total sleep time.

In a second main GAMLSS, with SWE as the dependent variable, and controlling for the same factors as above, we found a significant positive main effect of the MTsat value of the bmGM (**p=0.01, p_corr_=0.03)** and of age (**p=0.04, p_corr_= 0.12**) (**Table 2**, **Fig. 2B**). The GAMLSS also yielded an interaction between the bmGM MTsat value and age (**p =0.01, p_corr_= 0.03**). As for SOL, to gain insight in this interaction, we split our sample in 3 subsamples. We found significant negative association within older subsample (**p=0.006**) (**Table 3**, **Fig. 2D**), with higher MTsat related to lower SWE, suggesting that the overall significant change in the slope or direction of the association between SWE and MTsat is driven by the older individuals of our sample.

Importantly, the main GAMLSS using the other sleep parameters of interest as dependent variables (Sleep efficiency, REMS percentage, theta REMS power and No. of REMS arousals) were not significantly associated with the bmGM MTsat values (**Table 2**) suggesting that associations were specific to, or at least stronger for SOL and SWE.

Overall, these results show that increased MTsat values over the bmGM go along with enhanced sleep as reflected by a faster sleep onset and more intense NREM sleep. The association significantly changes from age 18 to 31 y, however. The younger individuals of our sample, aged 18 to 20 y, mirror the overall association with higher MTsat values associated with better sleep metrics while the association seem to progressively decrease or even revert in the intermediate subgroup, aged 21 to 23, and in the older subgroup, aged 24 to 31 y.

### Associations between sleep metrics and the qMRI myelin marker is present across the different brainstem compartments

We assessed the regional specificity of the detected associations for the bmGM compartment and assessed potential associations between SOL and SWE with MTsat value computed over the other 2 brainstem compartments yielded by the segmentation procedure. The separate GAMLSS analyses with SOL or SWE, as dependent variable, and MTsat value in brainstem reticulate grey matter (brGM) and periaqueductal grey matter and posterior hypothalamus (bpGMpH) (while including age, BMI, TST, TIV, MPM sequence and scanner) yielded the same statistical outputs than the analyses focusing on the bmGM compartment. The associations between SOL and SWE with MT values are thus observed in other 2 brainstem compartments (**Fig. 3A-D**; **Supplementary Fig. S2, Table S3**).

**Figure 3:**
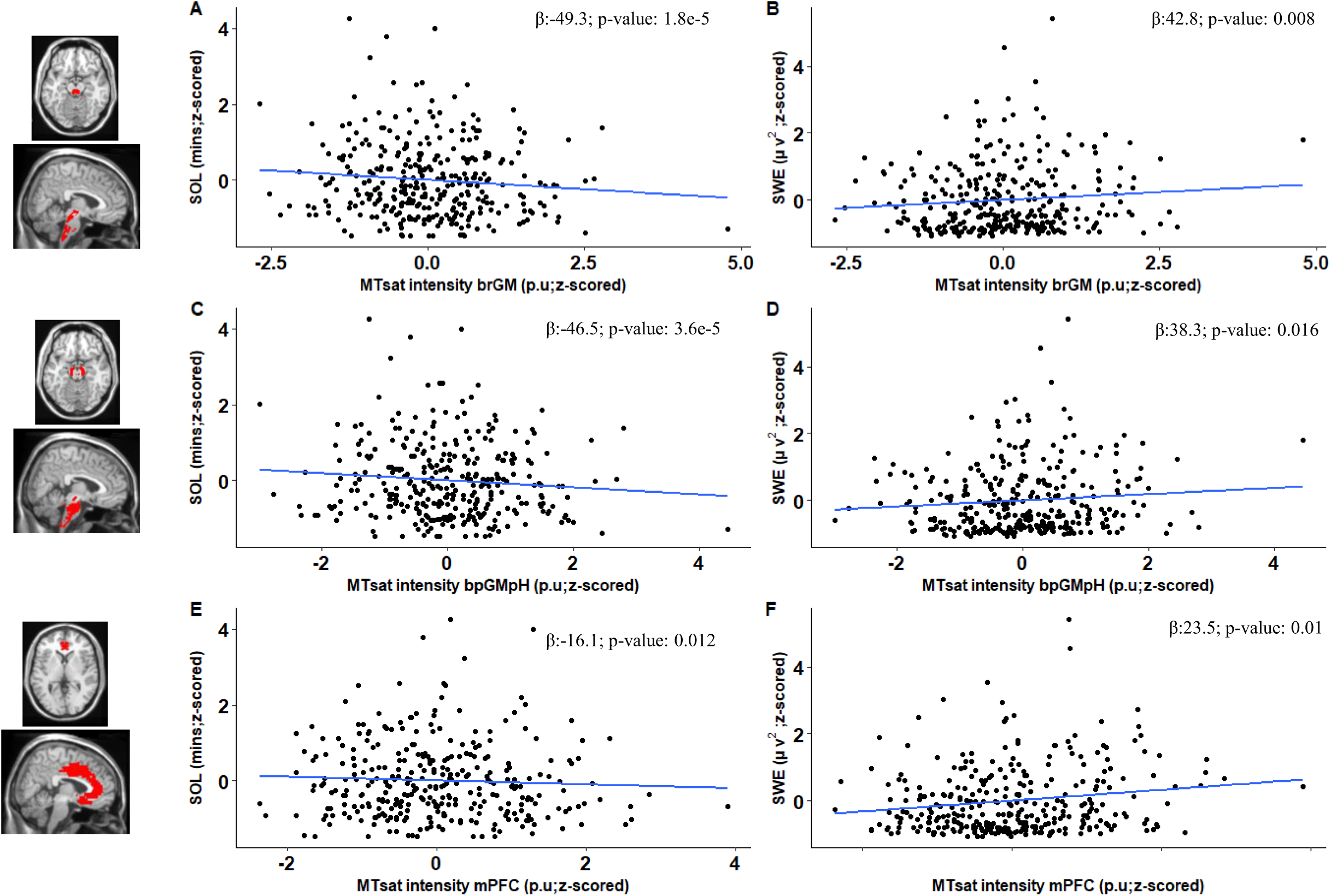
Plots (A-F) show association between sleep parameters and MTsat values for SOL and SWE with MTsat intensity in brainstem reticulate grey matter (brGM), periaqueductal grey matter and posterior hypothalamus (bpGMpH) and medial prefrontal cortex (mPFC), respectively, refer to Table S4 for statistical outputs of GAMLSSs.

### Associations between sleep latency and intensity go beyond associations with the qMRI myelin marker over the prefrontal cortex

We further assessed the regional specificity of the finding for the brainstem by considering the MTsat value computed over the medial prefrontal cortex (mPFC), which shows progressive myelination during adolescence and early adulthood and is highly sensitive to sleep homeostasis (McDougall et al., 2018, de Vivo and Bellesi, 2019). Separate GAMLSS revealed significant main effects of the MTsat value over the mPFC for SOL (**p= 0.01**; **p_corr_=0.03**) and SWE (**p=0.01**; **p_corr_=0.03**) (**Suppl. Table S3; Figure 3: E-F**) (while controlling for age, BMI, TST, TIV, MPM sequence and scanner). The GAMLSS using SOL and SWE as dependent variable yielded interactions between the mPFC MTsat value and age (respectively, p = 0.03 and p = 0.03; **Table S3; Figure S3**).

Importantly, when including the mPFC MTsat values in the GAMLSS, the associations between SOL and SWE and the MTsat value computed over the bmGM remained significant both as main effect (SOL: **p=2.9×10^-5^, p_corr_=1.7×10^-4^** ; SWE: **p=0.01, p_corr_=0.03**) as well as in interaction effect between MT value and age (SOL: **p=6.7×10^-5^, p_corr_=4.0×10^-4^** ; SWE: **p=0.01, p_corr_=0.03**) (**Suppl. Table S4)**. Therefore, when including the myelin marker over the prefrontal cortex, both SOL and SWE bear significant associations with MTsat value computed over the bmGM. As final validation of our main results, we computed cross-validation analyses by splitting dataset randomly into 70:30 train and test datasets which revealed that regressions between SOL or SWE and MTsat values computed over the bmGM remained significant in both trained and test dataset as compared to the when full dataset is used (**Suppl. Table S5**).

## Discussion

As part of the efforts to improve the understanding of the variability in healthy sleep, we tested whether qMRI, which informs about brain microstructural integrity (Weiskopf et al., 2013), would be correlated to canonical electrophysiological metrics of sleep, when estimated over the bmGM compartment which encompasses some of the most critical sleep-wake nuclei. We focussed on the MTsat metric because, in healthy tissue, it is a surrogate marker for myelin content (Lauleet al., 2007, Callaghan 2014). We find that SOL and SWE are significantly related to brainstem MTsat values, with values associated better sleep linked to higher MTsat values. Despite the limited age-range of our sample, the associations changed with age, with MTsat values appearing mostly associated with better sleep composition in the younger subsamples. These associations are not specific to a particular compartment of the brainstem as they were also found when considering the other brainstem GM compartments. The associations with the brainstem are, however, explaining a different part of the variance in SOL and SWE than the significant association we also find for both metrics with MTsat values computed over the mPFC. Finally, none of the other sleep metrics of interest, related to overall SWS-REMS organisation and to REMS fragmentation and intensity, were found to be significantly correlated to MTsat value over the brainstem, suggesting that associations are stronger for, or specific to SOL and SWE.

Myelin surrounds axons and has an intuitive impact on the crosstalk between distant brain regions reflected in the oscillations of the EEG (Nunez et al., 2015). Myelin is not only part of the white matter, but is also present in the grey matter, either at the bases of long axons or over short axons. Myelination can therefore also affect short-range neuronal connectivity and synchrony (Zatorre et al., 2012, Nunez et al., 2015). Myelination and synaptic pruning reflect a brain maturation process that progresses throughout childhood and adolescence and tail off through early adulthood, resulting in grey matter volume reduction and increase in white matter volume (Arain et al., 2013, Jamieson et al., 2020b). This maturation contributes in part to the important changes detected in sleep duration and quality taking place over the first 25 y of life (Hagenauer and Lee, 2013). There is for instance a significant reduction in the delta EEG frequency of SWS in adolescents that moves gradually with age from the posterior areas of the brain towards the anterior areas finishing at the PFC, a pattern that mirrors known brain myelination (Feinberg et al., 2011) and cannot be explained solely by a reductions in cortical thickness (Campbell and Feinberg, 2009).

Despite regional difference in myelin content across brainstem territories, an overall inverted U-shape reflect brainstem maturation with a progressive myelination up to ∼25y followed by a progressive decrease thereafter over the entire brainstem (Bouhrara et al., 2021). We find that several markers of sleep quality are associated with the brainstem myelin marker. While we expected specific correlations with the bmGM compartment, we find significant associations with all the GM compartments of the brainstem, the 3 GM. The fact that MTsat values over the brainstem and mPFC explain distinct parts of the variance in SOL and SWE, further support that regional maturation and integrity of the brainstem is important to some aspects of sleep.

The cross-sectional nature of our study does not allow inferring about causality. In addition, several nuclei of the brainstem are keys to sleep initiation and regulation and could contribute to the associations we detect. Other type of data (higher resolution, other contrasts, etc.) and/or the development of other segmentation tools [e.g. (Bazin et al., 2020)] are required to isolate specific nuclei. One can nevertheless posit that the association between MTsat and SOL or SWE are related to the maturation and integrity of the LC (though LC projection are mainly unmyelinated), raphe or cholinergic nucleus, which would optimize their local cross-talk - between neurons within and across nuclei – and their influence on cortical activity. This optimization would facilitate sleep onset and regulate slow wave production through and impact on the synchrony of neuron up and down states across patches of the brain. A relative silencing of the LC is for instance required to initiate sleep while transient noradrenalin release is locked to individual slow wave and spindles during SWS (Scammell et al., 2017, Osorio-Forero et al., 2022). Similarly, activity of the serotoninergic neurons of the dorsal raphe (DR) is high during wakefulness, decreases during NREM sleep, and ceases during REM sleep (Kato et al., 2022). Likewise, through their projection to the reticular nucleus of the thalamus, cholinergic neurons of the PPT and LDT facilitate sleep initiation and contribute to the production of spindle and therefore to the synchrony of cortical neuron activity (Xu et al., 2015, Ni et al., 2016).

*In vivo* studies of adolescent animal models showed that disruption of slow wave sleep (N3 stage) results in significant alterations in the development of brain connectivity (Jamieson et al., 2020a). Still in rodents, oligodendrocyte precursor cells (OPC), programmed to generate myelin sheath and GM and WM (Young et al., 2013, Yeung et al., 2014), proliferate two times faster during sleep than during wakefulness (Bellesi et al., 2013). Sleep is also involved in synaptic pruning and long-term potentiation and depression of newly form neurites (Puentes-Mestril et al., 2019), which may be wrapped in myelin sheets. The relationship between myelination and sleep has therefore been suggested to be bidirectional. In other words, our results are compatible with an impact of myelination of the diverse nuclei of the reticular formation on sleep quality as well as with an impact of sleep on myelination, where, for instance, repeated nights of sleep comprising more slow waves would increase myelin content. These hypotheses are not mutually exclusive and may take place concomitantly. Alternatively, our finding may also be incidental and arise from phenomenon that would both affect sleep and myelin. This phenomenon may also arguably be in part genetic which dictate brain myelin content and therefore affect sleep (Cirelli, 2009, de Vivo and Bellesi, 2019).

Importantly, while higher myelin content, as indexed by MTsat, may favor or be favored by better sleep quality in early adulthood, we find that the association may be reversed passed age ∼21 y. Myelination follows a temporally symmetric time course across the adult life span and exhibit an inverted U-shape association with age, with a peak at ∼25y, in several brainstem substructures similar to what is observed in cerebrum, although substantially less pronounced (Bouhrara et al., 2021) We posit that our findings could mean that delays the maturation of the brainstem across the ascending part of this inverted U-shape, i.e. over the second and third decade of life, is associated with lower sleep quality – with significant difference starting at ∼20-25y. Our interpretation is, however, based on significant interaction that we could partially draw from the significant association with specific age subgroups. Our findings are in line with the previously proposed occurrence of bi-directional interactions between sleep and myelin plasticity and endorse that sleep-wake-plasticity interactions may occur not only in age-specific manner but also in brain region-specific manner (Kurth et al., 2016). In line with this assumption, we recently reported in a whole-brain analysis of large part of the current sample that early night frontal SWA was negatively associated with regionally decreased myelin estimates, as indexed through MTsat values, in the temporal portion of the inferior longitudinal fasciculus (Deantoni et al., 2023). These results and the present ones show that regional myelin change may affect different aspects of sleep. A full appreciation of these region-specific associations over the lifespan requires more investigation, for instance using a sample with a larger age range of the healthy participants.

Our study bears limitations. We only included men and did not include a longitudinal component to the protocol. To circumscribe the multiple comparison issue, we also only focussed on six sleep metrics and a single qMRI parameter. In addition, MTsat values are not specific associated to myelin but rather to all macromolecules of the tissue. In healthy individuals, this macromolecular content mainly concerns cell membrane and therefore myelin of oligodendrocytes (Saccenti et al., 2020). MTsat may also be more directly related to the density of neurons and therefore indirectly related to myelin (Edwards et al., 2023). Other qMRI parameters depend to a lesser extent on myelin such as R2*, which is mainly driven by iron content (Stuber et al., 2014). It would be of high interest to consider other qMRI metrics along with MTsat as it would be to focus on other parts of the brain. Scarce studies reported that changes in the volumes of the hypothalamus and thalamus were associated to alterations in sleep, though mostly in clinical populations (narcolepsy, obstructive sleep apnea, or neurodegeneration) (Bartlett et al., 2019, Kreckova et al., 2019). Also, a reduction in thalamic GM density which could arguably result from synapses and myelin loss was reported to mediate part of the oscillatory changes in the EEG with ageing (Fitzroy et al., 2021).

Our study contributes to a better understanding of the brain correlates underlying sleep variability in healthy individuals. It may also have implication for patient populations that suffer from sleep disorders or degradation of myelin or both. For example, lesions at the brainstem and spinal cord level are more frequently associated with the appearance of REMS behaviour disorder (RBD) and restless legs syndrome (RLS) which constitute risk factors for Parkinson’s disease (Foschi et al., 2019). Likewise, sleep complaints are frequent in multiple sclerosis (Koltuniuk et al., 2022) and, based on our finding, myelin degradation over the brainstem could contribute to such complaints.

## Methods

### Ethics Statement

Approval for this study was obtained from the Ethics Committee of the Faculty of Medicine at the University of Liège, Belgium. Written informed consent was obtained from each participant prior to their participation and they were financially compensated.

### Participants

The study sample and study protocol is as previously published (Muto et al., 2021). We recruited a cohort comprising 364 healthy young men (aged 18-31) recruited between 2011 and 2015 as part of a large study assessing association between sleep architecture and circadian rhythmicity incorporating genetics and polygenic risk score assessments [for details see (Muto et al., 2021)]. The study only incorporated men to increase genetic homogeneity. The participants were excluded based on the following criteria: body mass index (BMI) > 27; presence of psychiatric history or severe brain injury; addiction; chronic medication affecting the central nervous system (CNS); smoking, excessive alcohol (> 14 units/week) or caffeine (>3 cups/day) intake; shift work in the past year; trans-meridian travels in the past 3 months; presence of moderate to severe subjective anxiety and depression as measured by the Beck Anxiety Inventory (BAI; score >16) (Beck et al., 1993) and Beck Depression Inventory II (BDI; score > 19) (Beck et al., 1988), respectively. Participants that exhibited poor sleep quality as assessed by the Pittsburgh Sleep Quality Index (PSQI) (Buysse et al., 1989) (score > 7), excessive daytime sleepiness as index by the Epworth Sleepiness Scale (Johns, 1991) (ESS; score >14) or excessive sleep apnea (apnea-hypopnea index > 15/h; 2017 American Academy of Sleep Medicine criteria, version 2.4) were excluded based on an in-lab screening night of polysomnography.

Out of 364, 28 participants were excluded after quality assessment of MRI data (12 had incomplete MRI acquisition, 15 had hyperintensity issues or movement artifacts in the MRI images, 1 had problem in segmentation process), 14 participants had incomplete baseline sleep electrophysiological data and 1 outlier resulting in a final sample of 321 participants. The characteristics of the final participant cohort are reported in **Table 1**.

### Sleep protocol

As described in [Muto 2021], individual sleep–wake history was strictly controlled: during the three weeks preceding the in-lab experiment, participants were instructed to follow a regular sleep schedule according to their habitual sleep timing (±30 min for the first 2 weeks; ±15 min for the last week; verified using actigraphy data - Actiwatch 4, CamNtech, Cambridge, UK). A urine drug test was performed (Multipanel Drug test, SureScreen Diagnostics Ltd) before completing an adaptation night at habitual sleep/wake schedule during which a full polysomnography was recorded to screen for sleep-related breathing disorders or periodic limb movements. On day 2, participants left the lab in the morning with the instruction not to nap which was confirmed with actigraphy data. They came back to the laboratory at the end of day 2 (3.5h hours before scheduled lights-off) and completed a baseline night of sleep (BAS) in full darkness centered on the average sleep midpoint of the preceding week. The current study focuses only on the baseline night of sleep. The remaining of the protocol included successively an extended nighttime sleep opportunity period, a daytime nap period, a normal night, a 40h sleep deprivation under constant routine condition (light < 5 lux) and finally a 12h recovery night. Outside sleep opportunity and sleep deprivation periods, participants were maintained in normal room light levels ranging between 50 and 1,000 lux depending on location and gaze.

### EEG acquisitions and analyses

Polysomnographic sleep data were acquired using Vamp amplifiers (Brain Products, Germany). The electrode montage consisted of 10 EEG channels (F3, Fz, F4, C3, Cz, C4, Pz, O1, O2, and A1; reference to right mastoid), 2 bipolar EOGs, 2 bipolar EMGs, and 2 bipolar ECGs. Acquisition on the screening night of sleep also included respiration belts, oximeter and nasal flow, 2 electrodes on one leg, but included only Fz, C3, Cz, Pz, Oz, and A1 channels. EEG data were re-referenced off-line to average mastoids. Scoring of sleep stages was performed using a validated automatic algorithm (ASEEGA, PHYSIP, Paris, France) in 30-s epochs (Berthomier et al., 2020) and according to 2017 American Academy of Sleep Medicine criteria, version 2.4. An automatic artefact and arousal detection algorithm with adapting thresholds (t Wallant et al., 2016) was further applied and artefact and arousal periods were excluded from subsequent analyses. Power spectrum was computed for each channel using a Fourier transform on successive 4-s bins, overlapping by 2-s, resulting in a 0.25 Hz frequency resolution. The night was divided into 30 min periods, from sleep onset, defined as the first NREM2 (N2) stage epoch, until lights-on. Averaged power was computed per 30 min bins, adjusting for the proportion of rejected data, and subsequently aggregated in a sum separately for REM and NREM sleep (Dijk and Landolt, 2019). Thus we computed slow wave energy (SWE) - cumulated power in the delta frequency band during N2 and N3 sleep stages, an accepted measure of sleep need (Dijk and Czeisler, 1995), and similar to that we computed the cumulated theta (4-8Hz) power in REM sleep. We then computed the cumulated power over the remaining EEG bands, separately for NREM and REM sleep: alpha (8-12Hz), sigma (12-16Hz), beta (16-25Hz) and theta (4-8Hz) bands. As the frontal regions are most sensitive to sleep–wake history (Cajochen et al., 1999), we considered only the frontal electrodes (mean over F3, Fz, and F4), as well as to facilitate interpretation of future large-scale studies using headband EEG, often restricted to frontal electrodes.

Our analyses focused on six sleep metrics to limit issues of multiple comparisons while spanning the most important aspects of sleep EEG: 1) sleep onset latency (SOL); 2) sleep efficiency (ratio sleep time – including N1 stage - vs. time in bed between light-off and light-on); 3) SWE during NREM sleep; 4) number of arousals during REM sleep; 5) REM sleep percentage and 6) cumulated theta power during REM sleep

### Image Acquisition and Quantitative Maps Creation

On average 3 weeks upon completion of the sleep study protocol, participants were examined on a 3T MR system. Out of the 335, neuroimaging data of 291 participants were acquired on a 3 T head-only MRI-scanner (Magnetom Allegra, Siemens, Erlangen, Germany). Further, out of these 291 participants, 191 were scanned using an Echo planar imaging (EPI) sequence, while 100 were scanned using an Actual Flip Angle Imaging (AFI) sequence. MRI data for the remaining 44 participants were acquired on a 3 T whole-body MRI-scanner (Magnetom Prisma, Siemens, Erlangen, Germany), with EPI sequence due to scanner replacement. The acquisition included a whole brain quantitative multiparameter protocol (MPM) as described in (Tabelow et al., 2019) and (Weiskopf et al., 2015).

The MPM protocol used in the present study has been already validated for multi-centric acquisitions (Weiskopf et al., 2013). Briefly, it consists of three co-localized series of 3D multi-echo fast low angle shot (FLASH) acquisitions at 1×1×1mm^3^ resolution and two additional calibration sequences to correct for inhomogeneities in the Radio Frequency (RF) transmit field (Lutti et al., 2012). The FLASH data sets were acquired with predominantly proton density (PD), T1, and magnetisation transfer (MT) weighting, referred to in the following as PDw, T1w and MTw echoes. Volumes were acquired in 176 sagittal slices using a 256 × 224 voxel matrix.

Quantitative multi-parametric volumes (PDw, T1w and MTw) were auto-reoriented against an MNI template and MT saturation (MTsat), PD, R1 and R2* maps were created using the hMRI toolbox (Tabelow et al., 2019) (http://hmri.info) implemented as an add-on toolbox for SPM12 (Statistical Para-metric Mapping, Wellcome Centre for Human Neuroimaging, London, UK, http://www.fil.ion.ucl.ac.uk/spm, University College London, revision 12.4). Here, we focused on the magnetization transfer saturation map (MT) which is related to the exchange of magnetization between mobile water protons that are bound to macromolecules as found in myelin. MT intensity values have been closely related to myelin content as shown in postmortem studies (Schmierer et al., 2004). Contrary to the commonly used MT ratio (percentage reduction in steady state signal), the MT saturation map explicitly accounts for spatially varying T1 relaxation time and flip angles (Helms et al., 2021).The map creation module includes the determination of B1 transmit bias field maps for transmit bias correction of the quantitative data. For this dataset, two different methods were used, the EPI (Lutti et al., 2012) and Actual Flip Angle Imaging (AFI) method (Yarnykh, 2007), according to the sequence implemented at the time of acquisition.

### Quantitative Multiparametric MRI Data Analysis

For the extraction of quantitative MTsat-derived myelin markers from specific brainstem regions MT and PD maps were first segmented into grey, white, and CSF tissue class maps using Unified Segmentation (US) within SPM12 (Ashburner and Friston, 2005). The whole-brain segmentation outputs were then diffeomorphically registered to a study-specific template, compatible with the MNI space, created using Shoot toolbox in SPM12 (Ashburner and Friston, 2011) in order to generate deformation fields that were used to warp MT and PD maps into the study-specific average space. Brainstem segmentation was then performed using unified segmentation with brainstem subregion tissue probability maps that were generated according to previously described methods based on a modified multivariate mixture of Gaussians (Lambert et al., 2013). Out of the 4 brainstem tissue classes produced, only tissue class 1 was considered for our main analysis, as it includes the substantia nigra, locus coeruleus and raphe nuclei designated as “brainstem monoaminergic grey matter,” (bmGM).The other two tissue classes mainly contained dorsal cranial nerve nuclei and were used for exploratory analyses only [tissue class 2: nucleus reticularis throughout its length and the pontine nuclei (reticulated grey matter; brGM); tissue class 3: Periaqueductal grey matter and posterior hypothalamus (bpGMpH) (Ridgway et al., 2009). The tissue class 4: brainstem white matter (bWM) was excluded from the analysis. The 3 tissue classes were warped back to individual space, using inverse deformation fields. Tissue class-specific smoothing (full width at half-maximum of 3 mm isotropic) was applied, and mean tissue class images were created after averaging in SPM Masking toolbox.

For MTsat-derived myelin markers from mPFC, segmented images were normalized using geodesic shooting (Ashburner, 2007) and smoothed with a tissue-weighted, for GM and WM separately, kernel of 4mm FWHM. Using the TD– ICBM Human atlas as implemented in the WFU-Pickatlas toolbox v 3.0.5b for SPM12 (Maldjian et al., 2003) we defined a mask the medial PFC via TD broadman areas in MNI space. For creating medial PFC mask, BA 24, BA 25 and BA 32 regions were merged. The modulated spatially normalised tissue maps of grey matter warped into MNI space were used for voxel-based quantification analyses (VBQ). Matlab based REX toolbox (https://web.mit.edu/swg/software.htm) was used to extract the mean MTsat intensity values (single-subject beta values) across each ROI mask.

### Statistical analysis

All analysis was carried out within R environment (version >4.1.3) (R Development Core Team, 2017). We employed generalized additive models for location scale and shape (GAMLSS) (Rigby and Stasinopoulos, 2005, Stasinopoulos and Rigby, 2008) to separately test the associations between six sleep metrics of interests (SOL, SWE, Sleep efficiency, REM percentage, REM power and No. of arousals in REM), as dependent variable, and the estimated MTsat values from bmGM as an independent variable. GAMLSS was selected based on data distribution and lowest AIC values among General Linear Models (GLM) and GAM models. In order to control for potential association between the sleep metrics and cortical MTsat values, average MTsat value was computed over the medial prefrontal cortex, as it is located below the electrodes of interest and an important source of slow waves (Nir et al., 2011). We further checked the specificity of associations with bmGM tissue class and computed MTsat intensity values from other brainstem tissue classes (brGM and bpGMpH) used as independent variable for subsequent analysis. bWM was not included in the analysis as it may be extremely heterogeneous, with motor (pyramidal) and somatosensory tracts. GAMLSS are univariate distributional regression models, where all the parameters of the assumed distribution for the dependent variable can be modelled as additive functions of the predictor/independent variables. On the other hand, GLMM and GAM are restricted to the exponential family of distributions (Rigby and Stasinopoulos, 2005). GAMLSS offer a large variety of distributions with up to four parameters - location (eg. Mean), scale (eg. variance), shape (eg. skewness) and shape (eg. kurtosis), classically noted as µ, σ, ν and τ. While only µ is modelled in (G)LM(M) and GAM(M), in GAMLSS all four parameters can be modelled, either with linear parametric, non-linear parametric or non-parametric (smooth) functions of the predictors. GAMLSS algorithms have been mainly designed to complete two tasks: maximize a penalized log-likelihood function addressing the estimates of fixed and random parameters, and evaluate the various smoothing parameters appropriately (Rigby and Stasinopoulos, 2014, Stasinopoulos et al., 2017, Rigby et al., 2019).

In the present study, either generalised Gamma (GA) or Gamma Lopatatsidis-Green (GG) family distribution was used for the GAMLSS regression modelling based on the Q-Q plot. Age, BMI, total sleep time (TST) and total intracranial volume (TIV) were included as covariates. Further, MPM sequence type and MRI scanner type was also included in the regression models to control for the protocol difference and scanner type during acquisitions. For age-subgroup interaction analysis GAM was used with either SWE or SOL as DV and age as categorical independent variable (IV) x MT value as continuous IV. Prior to the analysis, influential outliers were screened using worm plot. The worm plot (a detrended Q-Q plot) is a diagnostic tool for checking model fit, improving the fit and comparing the fit of different models (van Buuren and Fredriks, 2001). Cross-validation analyses were used for validating the models by splitting dataset randomly into 70:30 train and test datasets and regression performance was measured using the Root Mean Square Error (RMSE) metric. Because of the exploratory nature of the hypotheses, Benjamini & Hochberg False Discovery Rate (FDR) correction for 6 independent tests was used to test for significant associations.

Optimal sensitivity and power analyses in GAMLSS remain under investigation (Timmerman et al., 2021). We nevertheless computed a prior sensitivity analysis to get an indication of the minimum detectable effect size in our main analyses given our sample size. According to G*Power 3 (version 3.1.9.4) (Faul et al., 2007, Faul et al., 2009) taking into account a power of .8, an error rate α of .01 (corrected for 5 tests), a sample size of 321 allowed us to detect small effect sizes r > .24 (2-sided; absolute values; confidence interval: .13 – .34; R² > .06, R² confidence interval: .017 – .12) within a linear multiple regression framework including 2 tested predictor (MT value, age) and 5 other covariates (BMI, TST, TIV, MPM sequence, Scanner type).

## Acknowledgements

PT, MD, MVE, EK, FC, CS, CP and GV are/were supported by the Fonds de la Recherche Scientifique —FNRS-Belgium. The study was supported by the Wallonia-Brussels Federation (Actions de Recherche Concertées—ARC—09/14-03), WELBIO/Walloon Excellence in Life Sciences and Biotechnology Grant (WELBIOCR-2010-06E), FNRS-Belgium (FRS-FNRS, F.4513.17 & T.0242.19 &3.4516.11), Fondation Recherche Alzheimer (SAO-FRA 2019/0025), University of Liège (ULiège), Fondation Simone et Pierre Clerdent, European Regional Development Fund (Radiomed project), Fonds Léon Fredericq. D.J.D. is supported by the UK Dementia Research Institute (DRI). We acknowledge Christian Lambert for providing the scripts to perform the brainstem segmentation.

## Data availability

The data and analysis scripts supporting the results included in this manuscript are publicly available via the following open repository: https://gitlab.uliege.be/CyclotronResearchCentre/Public/xxx <u>(to be done following peer reviewing and upon acceptance for publication / and editor request)</u>. We used Matlab scripts for EEG and MRI data processing, while we used R studio for statistical analyses. Researchers willing to access the raw data should send a request to the corresponding author (GV). Data sharing will require evaluation of the request by the local Research Ethics Board and the signature of a data transfer agreement (DTA).

