## Supplemental Data for "*In vivo* marker of brainstem myelin is associated to quantitative sleep parameters in healthy young men"

**Supplementary Table S1:** Results derived from GAMLSS when testing for associations between sleep parameters and age.

|  | **Age** | **BMI** |
| --- | --- | --- |
| **SOL** |  |  |
| Estimate | 0.0221 | -0.0601 |
| p-value (95% CI) | 0.11 (-0.007, 0.051) | **1.6e-4** (-0.091, -0.029) |
| **SWE** |  |  |
| Estimate | -0.071 | 0.037 |
| p-value (95% CI) | **0.0002** (-0.105, -0.037) | 0.08 (-0.005,0.078) |
| **SE** |  |  |
| Estimate | -0.003 | 0.0012 |
| p-value (95% CI) | **0.002** (-0.005, -0.001) | 0.24 (-0.001, 0.003) |
| **REM%** |  |  |
| Estimate | 0.002 | 0.009 |
| p-value (95% CI) | 0.69 (-0.006,0.009) | **0.03** (0.001, 0.017) |
| **REM Theta Power** |  |  |
| Estimate | -0.047 | 0.035 |
| p-value (95% CI) | **0.001** (-0.074, -0.021) | **0.03** (0.004,0.066) |
| **REM Arousal** |  |  |
| Estimate | 0.033 | -0.020 |
| p-value (95% CI) | **0.001** (0.015, 0.051) | 0.07 (-0.041, 0.002) |
| **MTsat bmGM** |  |  |
| Estimate | -0.004 | 0.006 |
| p-value (95% CI) | 0.05 (-0.009, 2.7e-05) | **0.01** (0.001,0.011) |

**Table S2: Distribution of sleep parameters in age subgroups**

| **Sleep Features** | **Age group 18-20y**, N = 104*^1^* | **Age group 21-23y**, N = 133*^1^* | **Age group 24-31y**, N = 84*^1^* | **p-value***^2^* |
| --- | --- | --- | --- | --- |
| SOL (mins) | 12 (7, 20) | 15 (9, 23) | 14 (8, 23) | 0.14 |
| SWE (µV^2^) | 2705791  (1283248, 4767593) | 1567987  (685145, 3674841) | 1247139  (698626, 2668418) | **<0.001** |
| SE (%) | 94.9 (93.0, 96.6) | 94.1 (91.2, 95.5) | 93.5 (91.2, 95.6) | **0.001** |
| REMS % | 26.1 (23.4, 29.6) | 26.8 (23.5, 30.1) | 27.1 (24.2, 30.4) | 0.4 |
| REMS Theta Power (4-8 Hz) | 99012 (68140, 155066) | 81622 (43994, 141848) | 77748 (41761, 117621) | **0.012** |
| REMS Arousal (N) | 21 (15, 31) | 22 (15, 29) | 27 (20, 38) | **<0.001** |

^1^Median (IQR); ^2^Kruskal-Wallis rank sum test

SWE: slow wave energy; SOL: sleep onset latency; SE: sleep efficiency; REMS: rapid eye movement sleep

**Supplementary Table S3.** Results derived from GAMLSS when testing for associations between sleep onset latency (SOL) or slow wave energy (SWE) and MTsat values computed over the 2 other brainstem tissue classes and medial prefrontal cortex (mPFC) with age as interacting variable.

| **SOL** |  |  |  |  |  |  |  |  |
| --- | --- | --- | --- | --- | --- | --- | --- | --- |
|  | **MTsat** brGM | **Age*MTsat** brGM | **Scanner** | **Sequence** | **BMI** | **Age** | **TST** | **TIV** |
| **Estimate** | -49.3 | 2.15 | -0.12 | 0.18 | -0.06 | -0.51 | 0.002 | 0.0005 |
| **p-value**  **(95% CI)** | 1.8e-5  (-62.7, -35.9) | 4.4e-5  (1.54, 2.76) | 0.32  (-0.37,0.12) | 0.03  (0.017,0.34) | 0.0003  (-0.09, -0.03) | 1.1e-4  (-0.66, -0.36) | 0.03  (0.0002,0.004) | 0.11  (-0.00012,0.001) |
|  | **MTsat** bpGMpH | **Age*MTsat** bpGMpH | **Scanner** | **Sequence** | **BMI** | **Age** | **TST** | **TIV** |
| **Estimate** | -46.5 | 2.02 | -0.14 | 0.18 | -0.06 | -0.52 | 0.002 | 0.0005 |
| **p-value**  **(95% CI)** | 3.6e-5  (-59, -34) | 8.3e-5  (1.45,2.59) | 0.28  (-0.38,0.11) | 0.026  (0.02,0.35) | 2.3e-4  (-0.09, -0.03) | 1.9e-4  (-0.67, -0.37) | 0.032  (2.2e-4, 3.6e-3) | 0.10  (-9.9e-5,1.2e-3) |
|  | **MTsat** mPFC | **Age*MT**sat mPFC | **Scanner** | **Sequence** | **BMI** | **Age** | **TST** | **TIV** |
| **Estimate** | -16.1 | 0.62 | -0.08 | 0.16 | -0.06 | -0.28 | 0.002 | 0.001 |
| **p-value**  **(95% CI)** | 0.012  (-24.6, -7.66) | 0.030  (0.24,1) | 0.49  (-0.32, 0.16) | 0.053  (-0.003, 0.33) | 0.0002  (-0.09, -0.028) | 0.047  (-0.47, -0.09) | 0.015  (0.0004,0.004) | 0.016  (0.0002, 0.002) |
| **SWE** |  |  |  |  |  |  |  |  |
|  | **MTsat** brGM | **Age*MTsat** brGM | **Scanner** | **Sequence** | **BMI** | **Age** | **TST** | **TIV** |
| **Estimate** | 42.8 | -1.91 | 0.03 | -0.14 | 0.034 | 0.4 | -0.001 | -0.0002 |
| **p-value**  **(95% CI)** | 0.008 (10.1,75.6) | 0.01  (-3.4, -0.42) | 0.87  (-0.33,0.39) | 0.24  (-0.36,0.09) | 0.12  (-0.01,0.08) | 0.03  (0.03,0.77) | 0.26  (-0.004,0.001) | 0.66  (-2.5e-4, -1.5e-4) |
|  | **MTsat** bpGMpH | **Age*MTsat** bpGMpH | **Scanner** | **Sequence** | **BMI** | **Age** | **TST** | **TIV** |
| **Estimate** | 38.3 | -1.71 | 0.028 | -0.19 | 0.04 | 0.38 | -0.001 | -0.0002 |
| **p-value**  **(95% CI)** | 0.016 (6.79,69.8) | 0.018  (-3.15, -0.26) | 0.88  (-0.35,0.41) | 0.23  (-0.36,0.09) | 0.11  (-0.01,0.08) | 0.05  (-0.01,0.77) | 0.26  (-0.004,0.001) | 0.64  (-0.0003, -0.0002) |
|  | **MTsat** mPFC | **Age*MT**sat mPFC | **Scanner** | **Sequence** | **BMI** | **Age** | **TST** | **TIV** |
| **Estimate** | 23.5 | -0.85 | 0.03 | -0.11 | 0.04 | 0.35 | -0.002 | -0.002 |
| **p-value**  **(95% CI)** | 0.01  (6.8,40.2) | 0.03  (-1.6, -0.099) | 0.85  (-0.3,0.37) | 0.34  (-0.33,0.113) | 0.0729  (-0.003,0.08) | 0.07  (-0.02,0.72) | 0.12  (-0.004,0.0004) | 0.015  (-0.002, -0.002) |

BMI: body mass index; mPFC: medial prefrontal cortex, brGM: brainstem reticulate gray matter compartment; bpGMpH: periaqueductal grey matter and posterior hypothalamus, MPM sequence: type of multiparameter sequence, 2 sequence types were used (N = 222 for type 1; N = 99 for type 2- see methods); MRI scanner: data were acquired on 2 separate scanners (N = 286 for scanner 1; N = 35 for scanner 2 – see methods); MTsat: magnetisation transfer saturation; SOL: sleep onset latency; SWE: slow wave energy; TIV: total intracranial volume; TST: total sleep time.

**Table S4:** Results derived from GAMLSS when testing for associations between sleep onset latency (SOL) or slow wave energy (SWE) and MTsat values computed over all three brainstem tissue classes with age as interacting variable and MTsat values computed over the medial prefrontal cortex (mPFC) as covariate.

| **SOL** |  |  |  |  |  |  |  |  |  |
| --- | --- | --- | --- | --- | --- | --- | --- | --- | --- |
|  | **MTsat bmGM** | **Age*MTsat bmGM** | **MTsat mPFC** | **Scanner** | **Sequence** | **BMI** | **Age** | **TST** | **TIV** |
| **Estimate** | -47.5 | 2.07 | -2.36 | -0.12 | 0.16 | -0.061 | -0.49 | 0.002 | 0.001 |
| **p-value**  **(95% CI)** | **2.9e-05**  (-61.0, -34.0) | **6.7e-05**  (1.45,2.68) | **0.04**  (-4.59, -0.13) | 0.33  (-0.37,0.12) | **0.05**  (0.002,0.33) | 1.1e-04  (-0.1, -0.03) | **1.5e-04**  (-0.64, -0.34) | **0.04** (0.0002,0.004) | **0.01**  (0.0003,0.002) |
|  | **MTsat brGM** | **Age*MTsat brGM** | **MTsat mPFC** | **Scanner** | **Sequence** | **BMI** | **Age** | **TST** | **TIV** |
| **Estimate** | -48.1 | 2.09 | -2.32 | -0.117 | 0.161 | -0.06 | -0.50 | 0.002 | 0.001 |
| **p-value**  **(95% CI)** | **2.6e-05**  (-61.5, -34.7) | **6.1e-05**  (1.48, 2.7) | 0.04 (-4.56, -0.09) | 0.35  (-0.36,0.12) | 0.55  (-0.001,0.32) | 1.3e-04  (-0.09, -0.03) | 1.4e-04  (-0.650, -0.35) | 0.04  (0.0002,0.004) | **0.01**  (0.0003,0.002) |
|  | **MTsat bpGMpH** | **Age*MTsat bpGMpH** | **MTsat mPFC** | **Scanner** | **Sequence** | **BMI** | **Age** | **TST** | **TIV** |
| **Estimate** | -45.5 | 1.98 | -2.39 | -0.13 | 0.17 | -0.06 | -0.51 | 0.002 | 0.001 |
| **p-value**  **(95% CI)** | 4.7e-05  (-58, -33) | 1.1e-04  (1.41, 2.54) | 0.03  (-4.62, -0.167) | 0.30  (-0.38, 0.11) | 0.05  (0.004, 0.33) | 9.8e-05  (-0.09, -0.03) | 2.3e-04  (-0.66, -0.36) | 0.04  (0.0002,0.004) | 0.01  (0.0003, 0.002) |
| **SWE** |  |  |  |  |  |  |  |  |  |
|  | **MTsat bmGM** | **Age*MTsat bmGM** | **MTsat mPFC** | **Scanner** | **Sequence** | **BMI** | **Age** | **TST** | **TIV** |
| **Estimate** | 42.9 | -1.92 | 4.76 | 0.03 | -0.11 | 0.04 | 0.41 | -0.001 | -0.002 |
| **p-value**  **(95% CI)** | 0.007  (11.1,74.6) | 0.008  (-3.37, -0.473) | 0.002  (2.6,6.92) | 0.854  (-0.33,0.4) | 0.353  (-0.33,0.12) | 0.094  (-0.005,0.078) | 0.023  (0.05,0.76) | 0.228  (-0.004,  0.0009) | 0.016  (-0.002, -0.002) |
|  | **MTsat brGM** | **Age*MTsat brGM** | **MTsat mPFC** | **Scanner** | **Sequence** | **BMI** | **Age** | **TST** | **TIV** |
| **Estimate** | 44.1 | -1.98 | 4.75 | 0.03 | -0.11 | 0.04 | 0.42 | -0.001 | -0.002 |
| **p-value**  **(95% CI)** | 0.006  (12,76.3) | 0.007  (-3.44, -0.513) | 0.002  (2.6,6.91) | 0.855  (-0.326,0.39) | 0.359  (-0.328,0.118) | 0.097  (-0.006,0.078) | 0.019  (0.061,0.787) | 0.234  (-0.004,0.001) | 0.016  (-0.002, -0.002) |
|  | **MTsat bpGMpH** | **Age*MTsat bpGMpH** | **MTsat mPFC** | **Scanner** | **Sequence** | **BMI** | **Age** | **TST** | **TIV** |
| **Estimate** | 40.2 | -1.8 | 4.78 | 0.03 | -0.11 | 0.04 | 0.42 | -0.001 | -0.002 |
| **p-value**  **(95% CI)** | 0.01  (9.11,71.3) | 0.012  (-3.22, -0.38) | 0.0023  (2.62,6.94) | 0.875  (-0.34,0.4) | 0.36  (-0.33,0.12) | 0.088  (-0.005,0.079) | 0.03  (0.037,0.803) | 0.23  (-0.004,0.001) | 0.015  (-0.002, -0.002) |

BMI: body mass index; mPFC: medial prefrontal cortex, brGM: brainstem reticulate gray matter compartment; bpGMpH: periaqueductal grey matter and posterior hypothalamus, MPM sequence: type of multiparameter sequence, 2 sequence types were used (N = 222 for type 1; N = 99 for type 2- see methods); MRI scanner: data were acquired on 2 separate scanners (N = 286 for scanner 1; N = 35 for scanner 2 – see methods); MTsat: magnetisation transfer saturation; SE: sleep efficiency; SOL: sleep onset latency; SWE: slow wave energy; TIV: total intracranial volume; TST: total sleep time.

**Table S5: Cross-validation output after 70:30 split with GAMLSS analysis.**

| **SOL** | **Train dataset (70%)** | | **Test dataset (30%)** | |
| --- | --- | --- | --- | --- |
|  | **Estimate** | **p-value** | **Estimate** | **p-value** |
| MT value bmGM | -46.75 | **6.2e-07** | -72.58 | **3.6e-11** |
| MTsat mPFC | -2.61 | 0.081 | -2.18 | 0.22 |
| Age* MTsat bmGM | 1.99 | **1.7e-06** | 3.3 | **1.5e-10** |
|  | **Estimate** | **p-value** | **Estimate** | **p-value** |
| MT value bmGM | -47.41 | **4.6e-07** | -73.1 | **3.1e-11** |
| Age* MTsat bmGM | 2.03 | **1.2e-06** | 3.31 | **1.6e-10** |
| **SWE** | **Train dataset (70%)** | | **Test dataset (30%)** | |
|  | **Estimate** | **p-value** | **Estimate** | **p-value** |
| MT value bmGM | 46.13 | **0.026** | 70.13 | **0.018** |
| MTsat mPFC | 3.96 | **0.005** | 8.36 | **4.3e-6** |
| Age* MTsat bmGM | -1.98 | **0.033** | -3.66 | **0.009** |
|  | **Estimate** | **p-value** | **Estimate** | **p-value** |
| MT value bmGM | 44.71 | **0.034** | 66.40 | **0.032** |
| Age* MTsat bmGM | -1.92 | **0.042** | -3.39 | **0.021** |

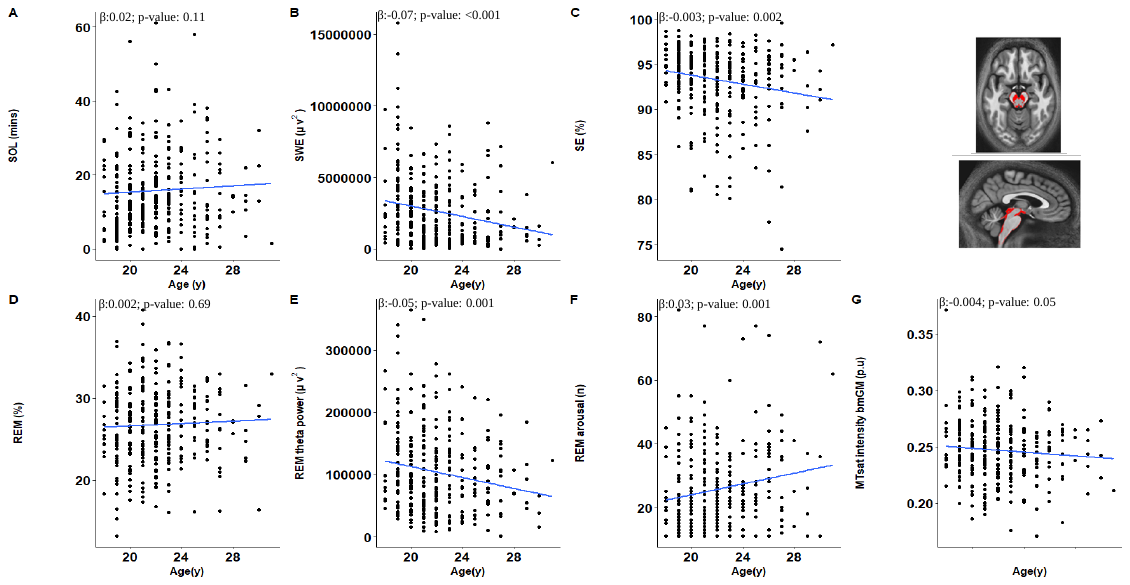

**Figure S1:** Plots (A-G) show results derived from GAMLSS when testing for associations between sleep parameters and age adjusted for BMI respectively, refer to Table S1 for statistical outputs of GAMLSSs.

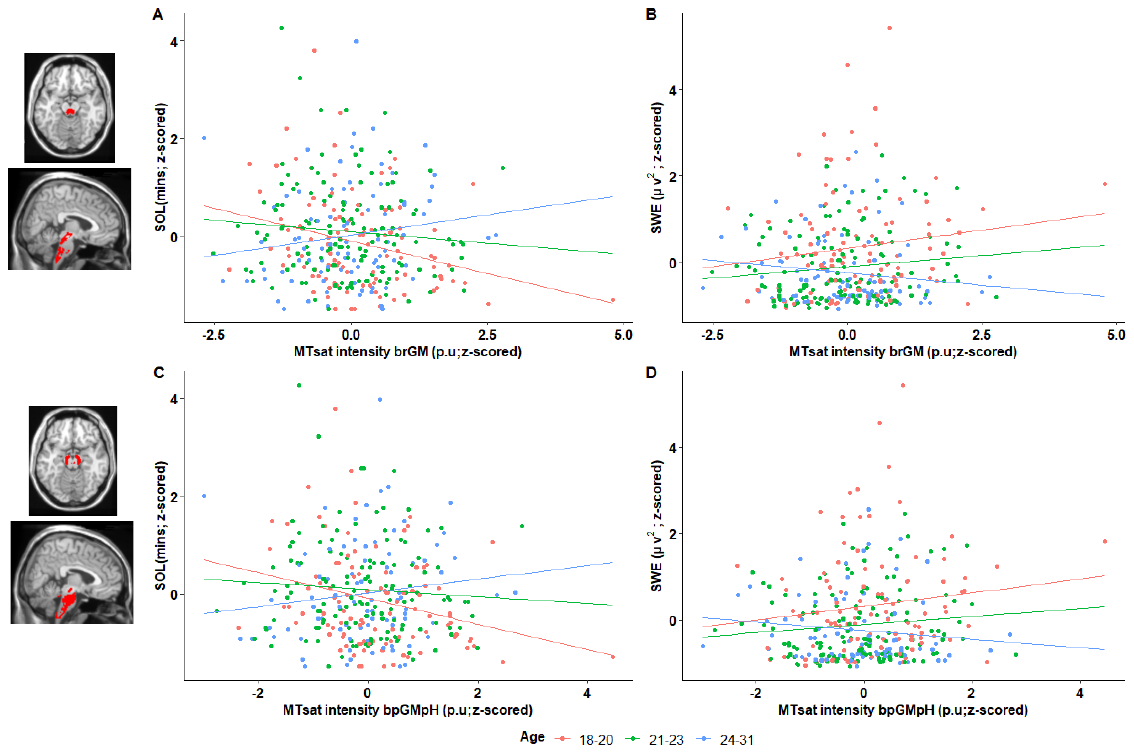

**Figure S2:** Plots (A-D) show association between sleep parameters and MTsat values by age group for SOL and SWE with MTsat intensity in brainstem reticulate grey matter (brGM), periaqueductal grey matter and posterior hypothalamus (bpGMpH) respectively.

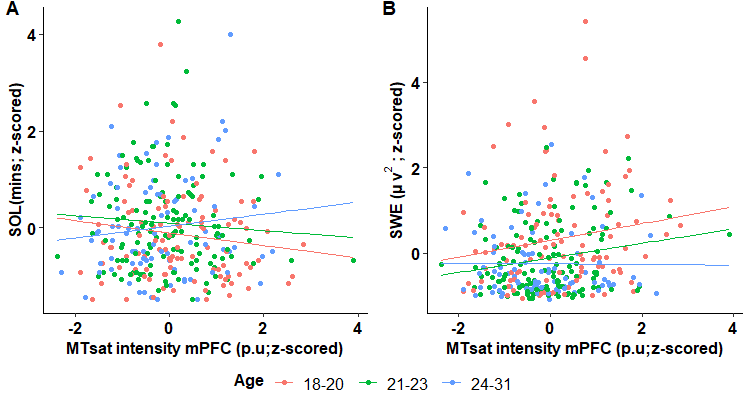

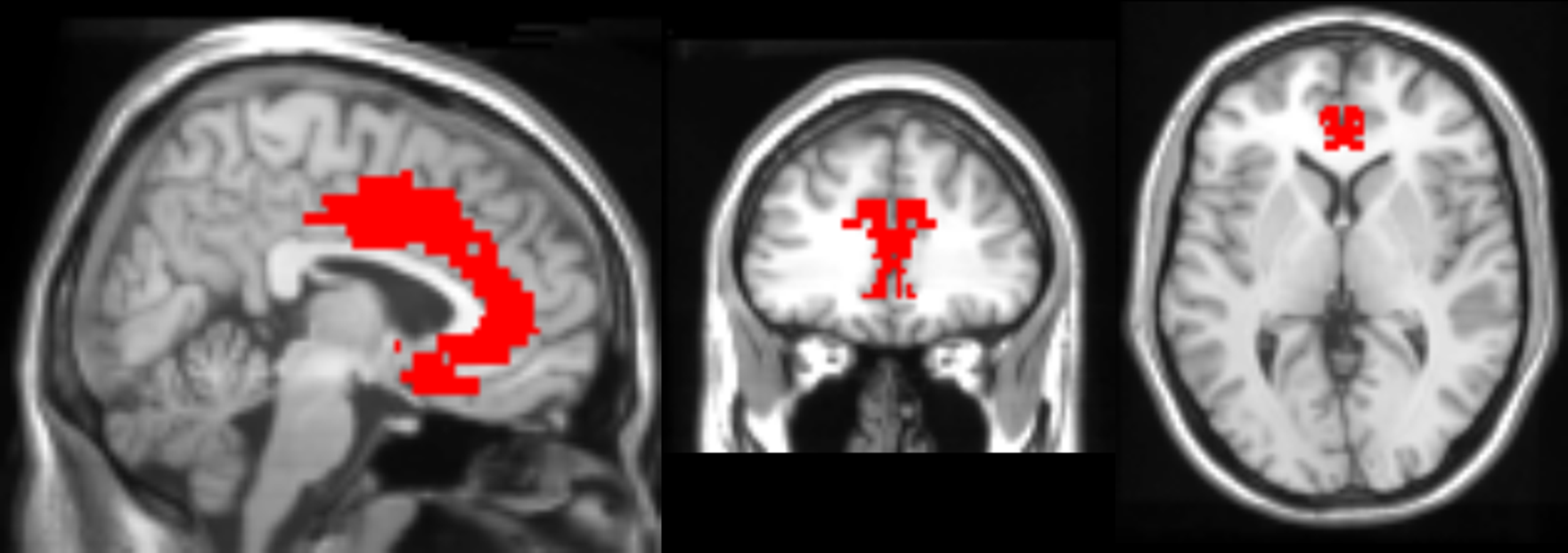

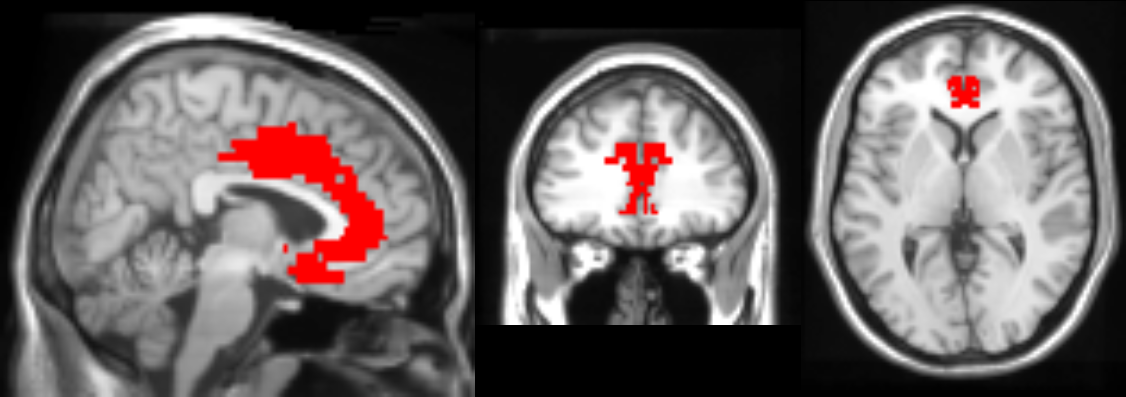

**Figure S3:** Plots (A-D) show association between sleep parameters and MTsat values by age group for SOL and SWE with MTsat intensity in medial prefrontal cortex (mPFC), periaqueductal grey matter and posterior hypothalamus (bpGMpH) respectively.
